# Fluid shear stress has a biphasic effect on clathrin-mediated endocytosis in vascular endothelial cells

**DOI:** 10.64898/2026.06.23.734037

**Authors:** Jie Yuan, Tomasz J Nawara, Leslie Donoghue Seeley, Yen T B Tran, Alexa L Mattheyses

## Abstract

Vascular endothelial cells (ECs) form a monolayer lining blood vessels and serve as a barrier between blood and tissues. Clathrin-mediated endocytosis (CME) is a major internalization pathway that involves a physical conformational change of the plasma membrane to form a vesicle and is therefore sensitive to the local environment. ECs are subjected to a myriad of fluid shear stress (FSS) rates from circulating blood, which we hypothesize affects CME. To test this, we used simultaneous two-wavelength axial ratiometry (STAR) microscopy, which provides nanoscale axial resolution, to determine the frequency and morphology of clathrin-coated vesicles as they form. Human umbilical vein endothelial cells (HUVECs) were transfected with dual-tagged clathrin light chain a (CLCa-iRFP-EGFP) and cultured under 10 dyn/cm^2^ FSS. CME activity was elevated in cells cultured under flow and assayed in static or flow conditions compared to statically cultured and imaged controls, indicating that FSS-induced changes to CME were maintained shortly after flow cessation. Single vesicle analysis showed cells cultured in FSS had a slight preference for vesicle formation with a flat-to-curved clathrin transition compared to control. Next, to assess the impact of different FSS rates, we cultured HUVECs at 20 and 40 dyn/cm^2^ FSS. We found total CME frequency was elevated compared to control at 20 dyn/cm^2^, but not 40 dyn/cm^2^. HUVECs cultured at both 20 and 40 dyn/cm^2^ had vesicles with increased lifetime and enhanced stability, as well as a higher proportion of vesicles formed through a flat-to-curved transition of clathrin.

## Introduction

Vascular endothelial cells (ECs) form a monolayer lining the inner surface of blood vessels, creating a tight barrier between blood and tissues and providing selective permeability across the vessel wall (Aghajanian et al., 2008; Kruger-Genge et al., 2019). In eukaryotic cells, clathrin-mediated endocytosis (CME) is a major pathway for internalizing macromolecules and membrane receptors. ECs internalize growth factors through paracrine signaling, including vascular endothelial growth factor (VEGF), which is critical for “stalk” and “tip” cell differentiation during angiogenesis, and fibroblast growth factor 2 (FGF-2), key for proliferation and migration of endothelial cells (Auciello et al., 2013; Genet et al., 2019). Other important cargo internalized via CME by ECs includes the adhesion molecule vascular endothelial cadherin (VE-cadherin) (Pitulescu and Adams, 2014; Xiao et al., 2005), soluble low-density lipoprotein (LDL) (Islam et al., 2022), transferrin (Mayle et al., 2012), and angiopoietin-1 (Ang-1) (Bogdanovic et al., 2009).

CME can be initiated by receptor-cargo binding, which recruits the coat protein clathrin via adaptor proteins such as AP2, forming clathrin-coated structures (CCS) at the plasma membrane (Mettlen et al., 2018). As clathrin is recruited, the nascent vesicle develops into an ‘Ω’-shaped dome. Finally, a clathrin-coated vesicle is released into the cytosol after scission mediated by the GTPase dynamin (Song et al., 2004). After internalization, many vesicles are destined for the endosome, where cargo is sorted for degradation or recycling back to the plasma membrane (Lakadamyali et al., 2006; Raiborg et al., 2001). Some internalized receptors continue signaling from endosomes, including epidermal growth factor receptor (EGFR), some G protein–coupled receptors (GPCRs), and transforming growth factor-beta receptor (TGF-βR) (Bakker et al., 2017; Balbis and Posner, 2010; Chen, 2009; Hanyaloglu and von Zastrow, 2008; Irannejad and von Zastrow, 2014; Seto et al., 2002).

Clathrin forms a triskelion structure in which the C-terminal domains of three clathrin heavy chains trimerize at a central hub, and the N-terminal domains extend outward with radial symmetry (Kirchhausen, 2000). The helix-rich N-terminal legs of the clathrin triskelion confer flexibility, enabling it to form a hexagonal meshwork on a flat membrane or a mixed hexagonal and pentagonal meshwork on a curved membrane (Kirchhausen, 2009; Royle, 2006). This flexibility allows flat clathrin lattices (FCLs) to convert into curved clathrin structures during vesicle formation (Chen and Schmid, 2020; McMahon and Boucrot, 2011). Two models have been proposed to describe clathrin-coated vesicle formation. The first is the constant curvature model, in which clathrin recruitment to the plasma membrane occurs concurrently with membrane bending. In this model, the vesicle radius remains relatively constant throughout vesicle formation. The second is the flat-to-curved model, also known as the constant area model, in which clathrin initially accumulates in a flat lattice. Subsequently, the clathrin-coated membrane transitions from flat to curved and then forms a vesicle (Mund et al., 2023; Obashi et al., 2023; Sochacki et al., 2021). A potential checkpoint mechanism that varies across different cargoes has been reported to affect membrane and clathrin curvature (Loerke et al., 2009; Mettlen et al., 2010; Mettlen et al., 2009). This flexibility might allow CME to accommodate internalization of different cargo sizes and types.

Formation of vesicles requires bending the plasma membrane, which is energetically unfavorable. Environmental changes that alter membrane tension, such as compression and osmotic pressure, have been shown to impact the frequency or dynamics of clathrin-coated vesicle formation (Ferguson et al., 2017). The dynamics of CME can also be modified from inside the cell. Manipulation of actin branching promoted initial vesicle invagination, favoring the constant curvature model (Yuan et al., 2026). Vascular ECs are heterogeneous, with differences based on position in the vascular tree (Jambusaria et al., 2020; Wakabayashi and Naito, 2023). Specialized subpopulations of ECs can be found within the capillaries of a single organ. Vascular ECs also experience varying degrees of pressure and shear force from blood flow, as well as mechanical forces from vasodilation and vasoconstriction. The mean fluid shear stress (FSS) in blood vessels is highest in arterioles (60 to 80 dyn/cm^2^), moderate in venules (20 to 40 dyn/cm^2^), and lowest in veins (< 1 dyn/cm^2^) (Ballermann et al., 1998). Vessel architecture, including curvature and bifurcation, as well as physical states such as exercise and meditation, also impact FSS (Weydahl and Moore, 2001; Zhao et al., 2000). Different flow environments promote transcriptional and morphological changes, including to the actin cytoskeleton (McCue et al., 2004; van der Meer et al., 2010). This raises the question of whether flow changes CME in ECs.

We previously found that cultivating HUVECs under 10 dyn/cm^2^ FSS promoted CME activity and contributed to vesicle formation through the flat-to-curved transition, compared with static controls (Nawara et al., 2025). In the present work, we build on this finding. First, we demonstrate that the flow-induced increase in CME is sustained in cells imaged under FSS and is therefore not due to membrane relaxation following the transition to a static environment. Next, we show that the change in CME rates is biphasic, which is higher than the static baseline at 10 and 20 dyn/cm^2^ and dropping to baseline at 40 dyn/cm^2^. Our data suggest that areas of the vasculature exposed to higher shear stress may have poorer cargo uptake compared to those exposed to lower shear stress, which could have proportionally much more uptake. The dynamics in vessels exposed to no flow, for example after stroke, may have limited drug uptake because of both lack of perfusion and a potential reduction in CME dynamics. Overall, this work indicates the importance of environmental factors in CME, and that the local environment should be considered when studying receptor-mediated signaling and drug uptake.

## RESULTS

### Increased CME persists in HUVECs after removal of flow

We previously found increased CME activity in HUVECs cultured under 10 dyn/cm^2^ FSS (Nawara et al., 2025). However, those measurements were made after transferring the cells from flow to static imaging conditions. The increase in internalization may have been due to FSS culture conditions, relaxation of the plasma membrane, or other changes caused by the environmental shift.

To test whether the transition from flow to static conditions contributed to changes in CME activity and vesicle formation dynamics, we cultured HUVECs in static conditions or 10 dyn/cm^2^ FSS generated by a peristaltic pump. After 24 hours, cells cultured in FSS exhibited a distinct morphology, with cell bodies elongated along the direction of flow (Fig. 1A). Next, we established 3 experimental conditions: 1) control cells cultured and assayed in static culture (S+S); 2) cells cultured under 10 dyn/cm^2^ FSS and assayed in static conditions (F+S); and 3) cells cultured and assayed under 10 dyn/cm^2^ FSS (F+F) (Fig. 1B). First, we performed a transferrin uptake experiment to test whether overall CME differed among groups. Cells cultured under 10 dyn/cm^2^ FSS had increased CME activity compared to control cells cultured in static conditions, regardless of whether they were assayed under flow or static conditions (Fig. 1C). Although transferrin concentration was the same across all samples, cells assayed under flow were exposed to a larger volume of media, resulting in greater total transferrin availability. To test whether more transferrin led to more uptake, we exposed statically cultured cells to twice the transferrin concentration and found no significant difference between the two static control groups (Supplemental Fig. 1). This suggests the increased transferrin internalization in HUVECs cultured and assayed under FSS was due to increased CME activity and not an increase in the overall amount of transferrin.

**Figure 1.**
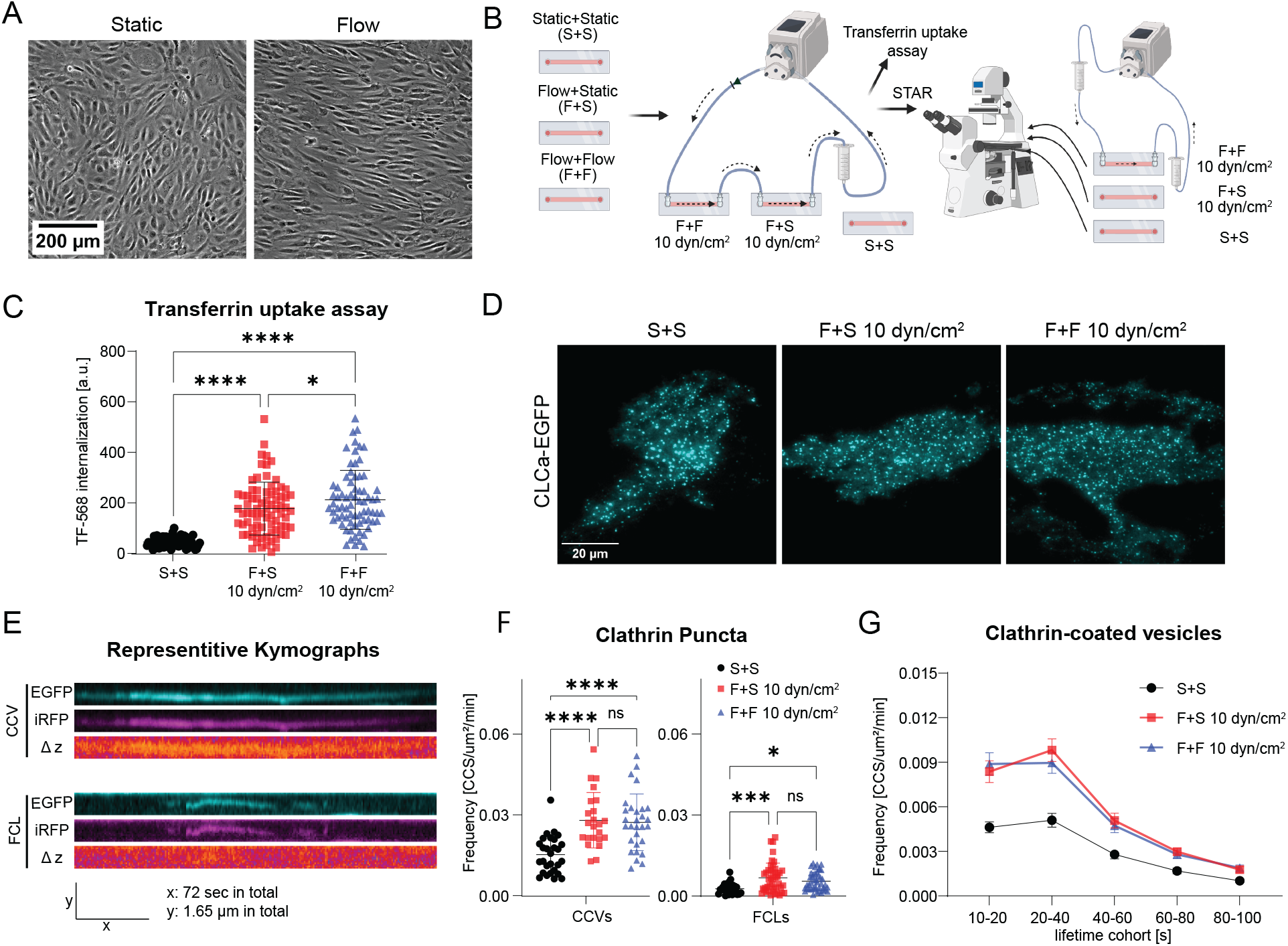
CME dynamics in HUVECs are increased by fluid sheer stress. (A) Phase contrast images of HUVECs cultured under static and flow (10 dyn/cm^2^) conditions. (B) Experimental design illustrating the experimental groups for STAR imaging and transferrin uptake. The groups are S+S, cultured and assayed in static conditions, F+S 10 dyn/cm^2^ cultured in fluid sheer stress (FSS) and assayed in static conditions, F+F 10 dyn/cm^2^ cultured and assayed in FSS conditions. (C) 20-minute transferrin uptake assay of HUVECs in control, group (F+S 10 dyn/cm^2^) and group (F+F 10 dyn/cm^2^) after 24 h static/flow culture. Cells for each group: control, 65 cells; F+S 10 dyn/cm^2^, 77 cells; F+F 10 dyn/cm^2^, 73 cells. Data was analyzed by one-way ANOVA with mean ± SD presented. (D) Example images of CLC-EGFP processed by DrSTAR from STAR microscopy for control, group (F+S 10 dyn/cm^2^) and group (F+F 10 dyn/cm^2^). (E) Kymographs of EGFP, iRFP and Δz signals from an individual clathrin-coated vesicle (CCV) and flat clathrin lattice (FCL). (F) Total density of CCVs and FCLs per μm^2^, per minute. Total cells from each group: S+S, 28; F+S 10 dyn/cm^2^, 23; F+F 10 dyn/cm^2^, 29; data from three independent experiments. Data were analyzed by one-way ANOVA separately for CCVs or FCLs, with mean ± SD presented. The same data set was used for Fig 1G. (G) Histogram of lifetime distribution of CCVs per μm^2^, per minute; mean ± SEM.

Next, we investigated whether single-vesicle formation dynamics were impacted by imaging under flow. To do this, we used STAR microscopy, which allows discrimination between curved clathrin vesicles and flat clathrin lattices (Nawara et al., 2022). However, the peristatic pump caused severe focus drift during high-resolution live-cell imaging. Therefore, we designed a gravity-driven flow system to provide constant 10 dyn/cm^2^ FSS during imaging (Fig. 1B). We transfected HUVECs with the dual-labeled STAR tag (CLCa-iRFP-EGFP), cultured them in static or flow conditions, and imaged them by STAR microscopy (Fig. 1B). Dynamic diffraction-limited clathrin puncta were observed in each group (Fig. 1D). The MATLAB tools DrSTAR and CMEAnalysis were used to process the data, identify dynamic clathrin puncta, and to distinguish curving clathrin-coated vesicles (CCVs) and flat clathrin lattices (FCLs) (Aguet et al., 2013; Nawara et al., 2023). Both CCVs and FCLs can result in increased intensity of EGFP and iRFP due to the accumulation of labeled clathrin. However, only CCVs showed a significant change in the ratio channel Δz reflecting their increased height relative to the plasma membrane as the vesicle is formed and clathrin curvature is established (Fig. 1E). The Δz channel will remain close to background for FCLs, allowing discrimination between these two dynamic clathrin populations (Fig. 1E) (Nawara et al., 2022).

Quantification of dynamic clathrin puncta revealed an overall increase in the frequency of both CCVs and FCLs in cells cultured under 10 dyn/cm^2^ FSS compared to control static-cultured cells (Fig. 1E, F). The total frequency of CCVs was not statistically different between the two flow-cultured groups, suggesting the change is driven by cellular changes during the 24 hours of culture under FSS, and not by FSS alone. This differed slightly from the transferrin uptake assay between these two groups (Fig. 1C), possibly because the transferrin assay measures total internalization, whereas STAR microscopy quantified CCVs only at the basal membrane.

To investigate the nanoscale dynamics of CME vesicle formation, we tracked the onset of membrane curvature, t(Δz), and the arrival of clathrin at the plasma membrane, t(CLC) (Fig. 2A). For each individual CCV, we calculated t(Δz) - t(CLC) and plotted the result as a cumulative frequency histogram for all CCVs. Both groups cultured under flow, whether imaged in static or flow conditions, showed a longer time delay between membrane curvature and clathrin arrival, indicated by a rightward shift of the curves, suggesting an increased flat-to-curved mechanism of vesicle formation (Fig. 2B). Next, we measured CCV lifetimes, defined as the interval from when the EGFP-CLC signal of CCV was first detected when it disappeared, across different groups and found culture in FSS extended CCV lifetime. The lifetime of CCVs did not differ between cells cultured under flow and imaged under flow or static conditions (Fig. 2C). Quadrant gating of lifetime vs [t(Δz) - t(CLC)] and lifetime vs mean squared displacement further confirmed that increased lifetime, a shift toward the FTC model, and reduced CCV stability occurred together when cultured under 10 dyn/cm^2^ FSS (Fig. 2D-F). These results showed that 10 dyn/cm^2^ FSS promoted CME activity and increased the lifetime and stability of CCVs. This effect persisted after cells were removed from flow.

**Figure 2.**
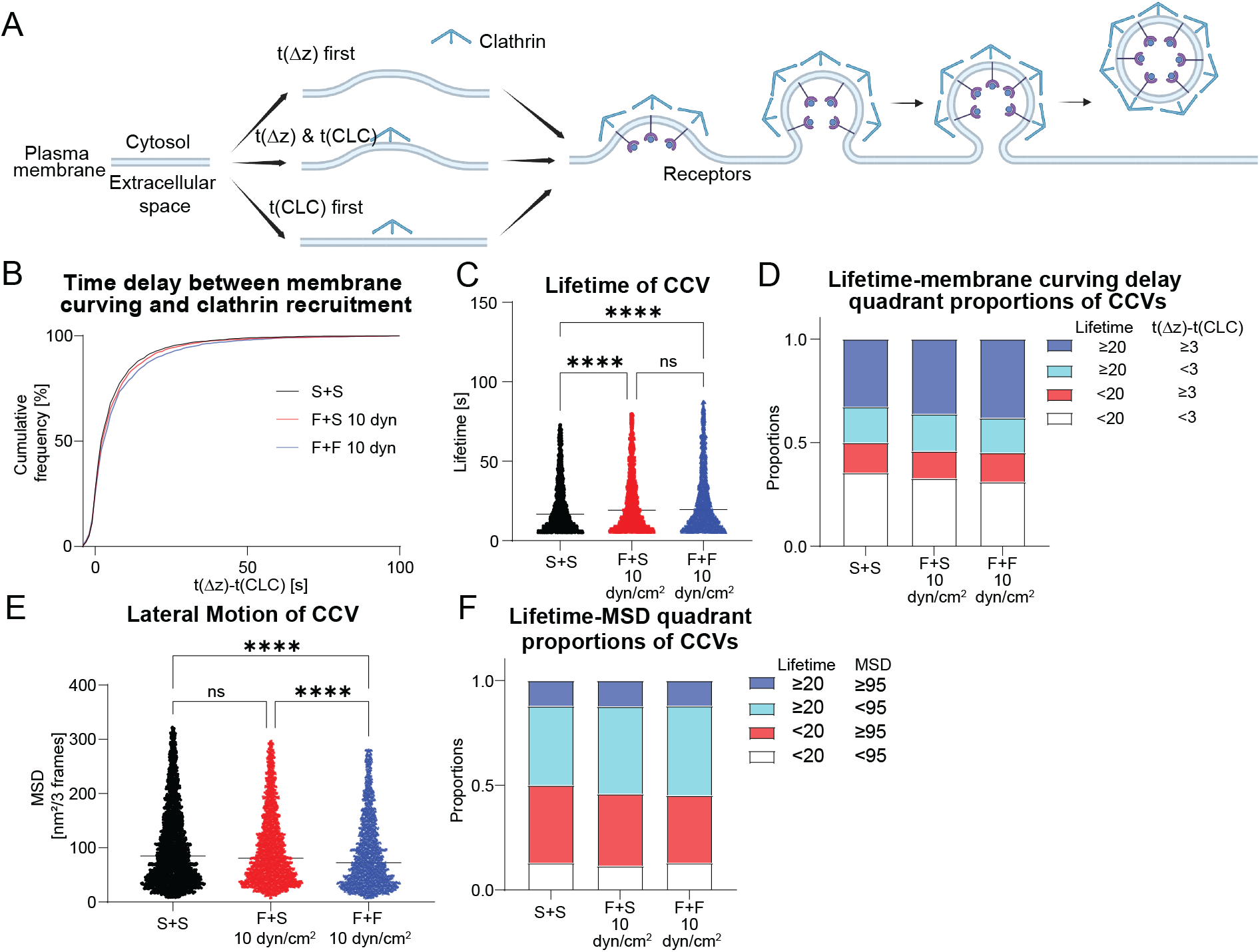
The flow-altered CME profiles in HUVECs remained shortly after cessation of flow. (A)Illustration for detection of membrane curving and recruitment of clathrin by STAR microcopy.(B)Cumulative histogram of the time between the initiation of membrane curvature and recruitment of clathrin. CCVs from each group: S+S, 3097; F+S 10 dyn/cm^2^, 3250; F+F dyn/cm^2^, 3442. The same data set was used for fig 2c to 2f. (C) Lifetime of CCVs with Kruskal-Wallis test. Median bar presented. Each data point represents a CCV. (D) Proportional distribution of CCVs by median quadrant of lifetime and membrane curving delay. (E) Motion analysis of CCVs and FCLs with a gap of three frames (0.9 s) by mean-square displacement (MSD) with Kruskal-Wallis test. Median bar presented. Each data point represents a CCV. (F) Proportional distribution of CCVs by median quadrant of lifetime and MSD.

### Flow rate has a biphasic effect on CME frequency

To test whether CME frequency would continue to increase at higher FSS, we cultured HUVECs at 20 and 40 dyn/cm^2^ FSS. Because we identified minimal differences in CME in cells imaged in static or flow conditions, HUVECs were imaged under static conditions by STAR microscopy (Fig. 3A). CLC-EGFP puncta were visible and dynamic in each group (Fig. 3B). At 20 dyn/cm^2^, overall CME activity was higher than the static group (Fig. 3C). However, when the FSS increased to 40 dyn/cm^2^, the overall CCV frequency was unchanged compared to the control, due to a reduction in short-lifetime CCVs (Fig. 3C,D). When the mode of vesicle formation was examined, we found that higher FSS promoted the flat-to-curved model of CCV formation and increased the lifetime of CCVs during CME vesicle formation at both 20 and 40 dyn/cm^2^ FSS (Fig. 3E-G). These changes tended to occur together across all CCVs, as shown by quadrant gating analysis of lifetime vs [t(Δz)-t(CLC)] and lifetime vs mean square displacement (Fig. 3H,I)

**Figure 3.**
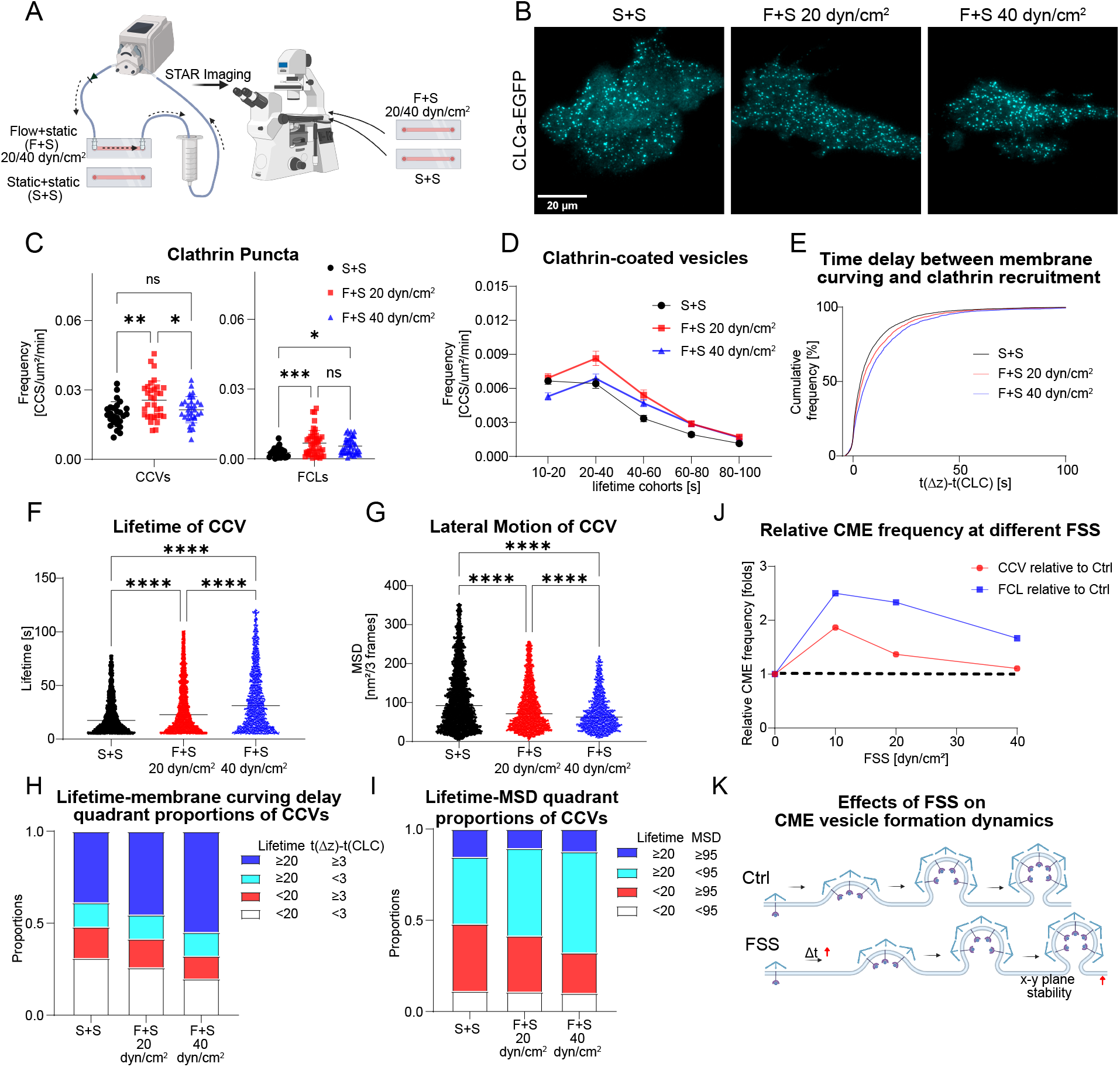
Increased flow rate has a biphasic effect on CME dynamics in HUVECs. (A) System of tracking CME dynamics in HUVECs at 20/40 dyn/cm^2^. (B) Example images of CLC-EGFP processed by DrSTAR from STAR microscopy for control, group (F+S 20 dyn/cm^2^) and group (F+S 40 dyn/cm^2^). (C) Total density of clathrin-coated vesicles (CCVs) and flat clathrin lattices (FCLs) per μm^2^, per minute. Total cells for three replicates from each group: S+S, 26; F+S 20 dyn/cm^2^, 33; F+S 40 dyn/cm^2^, 31. Data was analyzed by one-way ANOVA separately for CCVs or FCLs with mean ± SD presented. (D) Histogram of lifetime distribution of CCVs per μm^2^, per minute; mean ± SEM. (E) Cumulative histogram of the time between the initiation of membrane curvature and recruitment of clathrin. CCVs from each group: S+S, 2821; F+S 20 dyn/cm^2^, 2800; F+s 40 dyn/cm^2^, 1874. The same data set was used for Fig. 3F-I. (F) Lifetime of CCVs with Kruskal-Wallis test. Median bar presented. Each data point represents a CCV. (G) Motion analysis of CCVs and FCLs with a gap of three frames (0.9 s) by mean-square displacement (MSD) with Kruskal-Wallis test. Median bar presented. Each data point represents a CCV. (H) Proportional distribution of CCVs by median quadrant of lifetime and membrane curving delay. (I) Proportional distribution of CCVs by median quadrant of lifetime and MSD. (J) Relative CME frequency (In folds) over control group at different FSS. (K) Effects of FSS on CME vesicle formation dynamics.

## CONCLUSION AND DISCUSSION

In this work, we investigated how CME dynamics in HUVECs respond to varying shear stress. First, we found that CME frequency increased in HUVECs cultured under pulsatile flow at 10 dyn/cm^2^ FSS by the transferrin uptake assay, where internalization was assayed in both flow and static conditions. This increase of CME in HUVECs cultivated in FSS was also observed in our single-vesicle CME analysis where the frequency of CME was increased in cells imaged under both static and flow conditions.

Therefore, we continued to culture HUVECs at 20 and 40 dyn/cm^2^ and imaged the cells under static conditions. As FSS increased, the promoted CME dynamics became less distinct from those of the statically cultured group, as indicated by the relative CME frequency at different flow rates (Fig. 3J), which was calculated as the means of the experimental groups divided by the mean of the control groups for CCVs and FCLs (from Figs.1F and 3C). The stability and lifetime of CCVs increased during CME vesicle formation as FSS increased (Fig. 3K). The clathrin also tended to stay on the membrane for longer before curving into vesicles as FSS increased.

FSS has been reported to increase membrane tension (White and Frangos, 2007) and promote actin expression in endothelial cells (Malek and Izumo, 1996). In our previous work, we found that increased membrane tension reduced the frequency of CCVs and that actin stabilizes CCVs during formation (Yuan et al., 2026). In this work, as FSS increased to 20 and 40 dyn/cm^2^, there was a gradual decrease in short-lifetime CCVs (Fig. 3D). This was possibly due to the further enhanced actin dynamics induced by higher flow rate.

Therefore, we propose that as the FSS generated by pulsatile flow increased from 0 to 10 dyn/cm^2^, HUVECs increased actin expression and became more active in CME. As the FSS increased to 20 dyn/cm^2^, further enhancement of the actin network may be required. This could inhibit the formation of CCVs and thereby contribute to the reduction in internalization. The situation deteriorated at a higher FSS of 40 dyn/cm^2^ where more abundant actin may be required to support the HUVECs against higher FSS but also restrict CME vesicle formation. However, the increased actin along increased FSS keeps extending the lifetime of vesicles and increasing vesicle stability. In addition to endothelial cells, FSS has also been reported to promote CME in renal tubular epithelial cells (Lackner et al., 2024; Raghavan et al., 2014; Xu et al., 2020).

In this work, the frequency of the peristatic pump increased as the FSS increased and therefore may have also contributed to changes in the actin architecture and CME.

In summary, our work demonstrates that moderate levels of FSS promote CME activity in ECs, but this effect did not persist at higher flow rates, possibly due to an increased actin meshwork. This work provides insight into endothelial cell internalization under varying blood flow rates and underscores the need to study endocytosis under physiologically relevant environmental conditions.

## MATERIALS AND METHODS

### Cell culture and transfection

Human Umbilical Vein Endothelial Cells (HUVECs, Sigma, 200P-05N) were cultured in vascular endothelial growth media (VEGM), prepared by mixing Vascular Cell Basal Medium (ATCC, PCS100030) and Endothelial Cell Growth Kit-VEGF (ATCC, PCS100041).

For transfection, HUVECs were plated in a 6-well plate (Fisher Scientific, 08-772-1B) at 70,000 cells per well in 2.5 mL VEGM. After 24 hours, HUVECs were switched to Dulbecco’s modified Eagle’s medium (DMEM; Corning, 10013CV) with 2% fetal bovine serum (FBS, Gibco, 10438-026). For transfection, 2 µg of plasmid DNA and 2 µL of transfection reagent (Sigma, TransIT-2020, MIR540) were diluted in 250 µL PBS (Fisher Scientific, MT21030CV), mixed by pipetting, incubated at room temperature for 20 min, and then added to 1 well of HUVECs. After 4 hours, the media was changed to VEGM.

For FSS experiments, 125,000 HUVECs transfected with the STAR plasmid were seeded into a µ-Slide with a 400 µm high channel (Ibidi, 80177). The µ-Slide was gently flushed with 1 mL VEGM after 1 hour. After 8 hours, the µ-Slide was flushed again with 1 mL VEGM containing 0.5% penicillin-streptomycin (100 IU/mL, Gibco, 15070-063). On the second day, the µ-Slide was flushed again with 1 mL VEGM containing 1% penicillin-streptomycin. After 24 hours, the µ-Slide was connected to the flow system.

### Fluidic culture system

A µ-Slide was connected to the flow system, and a peristaltic pump (Kamoer Fluid Tech, Amazon, DIPump550-B253) was used to provide pulsatile flow 10, 20, or 40 dyn/cm^2^ FSS. The pump tube was set up on the pump roller and the media reservoir, a 10 mL syringe (BD, 302995), was connected to the inlet of the pumping tube with Nalgene− 50 Platinum-Cured Silicone Tubing (Thermo Fisher, 8060-0020). The outlet of the pumping tube was connected to a unidirectional valve. The valve was then connected to the inlet of the µ-Slide using an adapter (Ibidi, 10831). The outlet from the µ-Slide was then connected back to the media reservoir. An air filter, Millex−-GS Filter Unit (Sigma, SLGSM33SS), was added to the media reservoir for gas exchange.

For the 10 dyn/cm^2^ group, the F+S and F+F µ-Slides were connected in series to the pumping system to ensure a uniform flow rate across the samples for 24 hours. Before imaging, the F+S group was disconnected from the pumping system and incubated with VEGM containing 5 μM biliverdin (Cayman Chemical Company, 19257) for 30 minutes, then flushed with 1 mL of fresh VEGM and imaged. For the F+F group, the media in the pumping system was replaced with 8 mL of VEGM containing 5 μM biliverdin, and HUVECs were cultured for 30 min under flow. The µ-Slide was then flushed with 1 mL of fresh VEGM and connected to the gravity-driven fluidic imaging system. For the S+S control group, the media was changed to VEGM containing 5 μM biliverdin for 30 min, then to fresh VEGM for imaging.

For the 20 and 40 dyn/cm^2^ groups, the µ-Slides for the static control and 20 dyn/cm^2^ were prepared simultaneously. On the second day, the control group was incubated with VEGM containing 5 μM biliverdin for 30 min in static conditions and then imaged by STAR microscopy in static conditions. The flow group F+S 20 dyn/cm^2^ was treated with VEGM containing 5 μM biliverdin for 30 min in the pumping system. Then F+S 20 dyn/cm^2^ slide was disconnected and imaged. The control slide was re-connected to the pumping system and cultured under 40 dyn/cm^2^ as F+S 40 dyn/cm^2^ group for 24 hours. On the third day, the F+S 40 dyn/cm^2^ group was incubated with 5 μM biliverdin for 30 min in FSS and imaged by STAR microscopy in static conditions.

### Transferrin uptake assay

Four groups were set up: S+S, F+S 10 dyn/cm^2^, F+F 10 dyn/cm^2^, and S+S, TFx2. After 24 h of fluidic or static culture, all slides were starved for 30 min in FluoroBrite (Gibco A1896701) medium and then incubated in FluoroBrite containing Transferrin-Alexa Fluor 568 conjugate (TF-568, Thermo Fisher Scientific, T23365) for 20 min. The S+S, F+S 10 dyn/cm^2^ and F+F 10 dyn/cm^2^ groups were treated with 25 μg/mL TF-568. The S+S TFx2 group was treated with 50 μg/mL TF-568. The S+S, F+S 10 dyn/cm^2^ and S+S, TFx2 groups were incubated in static conditions while the F+F 10 dyn/cm^2^ group was incubated in FSS. After 20 min of incubation with TF-568, slides were placed on ice to stop internalization, washed with cold acid 3 times (0.2 M acetic acid (Fisher Chemical, A38S-212), 0.2 M NaCl (Fisher Scientific, BP358), pH 2.5) to remove surface-bound TF-568, flushed with DPBS (Corning, 21030CV) 3 times, fixed with 4% PFA (Electron Microscopy Sciences, 15710) on ice for 20 min, and finally washed with DPBS 3 times. Widefield images were taken with using a ET/mCh /TR filter cube (Chroma) and differential interference contrast to assess Tf uptake and cell boundaries.

### STAR Imaging system

A Nikon Ti-2 microscope was used for live-cell STAR imaging. The microscope has a 60×1.49-NA objective, manual total internal reflection fluorescence (TIRF) illuminator (Nikon, TI-LA-TIRF), 488-nm (Obis, 488-150 LS), and 647-nm (Obis, 1196627) excitation lasers, fiber coupling optics: fiber mount (Thorlabs, MBT621D), converging and directing the laser objective (Olympus, RMS10X), optical fiber (Thorlabs, P3-405BPM-FC-2), C-NSTORM QUAD 405/488/561/638-nm TIRF dichroic. A stage-top incubator (Tokai Hit, INUBG2SF-TIZB) maintained 37 °C and 5% CO_2_ for live-cell imaging.

An Optosplit III (Cairn Research) image splitter, equipped with ET525/50 m and ET705/72 m emission filters (Chroma) and T562lpxtr-UF2 and T640lpxtr-UF2 dichroic mirrors, was added between the microscope and the camera (Hamamatsu). When both 488-nm and 647-nm were excited, the fluorescence emissions from two fluorescent tags were split onto adjacent but distinct regions on the ORCA-Flash 4.0 v3 scientific complementary metal-oxide-semiconductor camera.

The system was coupled to a data acquisition device (NIDAQ, National Instruments, BNC-2115) and controlled using Nikon Elements software (version 5.02) and Coherent Connection software (version 3.0.0.8). The Optosplit III was calibrated according to the manufacturer’s protocol using the NanoGrid (Miraloma Tech, A00020). Images were acquired at 0.3 seconds per frame, with a 200 ms exposure time and simultaneous 488- and 647-nm excitation. The total imaging time for each cell was 5 min.

### Gravity-driven fluidic imaging system

To integrate flow system with the STAR microscope, a gravity-flow system was devised. A 0.75-meter-high post (Thorlabs, PSY321) was mounted on the optical table (Thorlabs, SDA90120) with two aluminum breadboards (Thorlabs, MB12, PSY321/S). The upper breadboard held a 10 mL syringe (BD, 302995) with silicone tubing (Thermo Fisher, 8060-0020) descending to the stage-top incubator. The tubing wrapped around the water bath inside the incubator before connecting to the µ-Slide inlet. The µ-Slide outlet was connected to a reservoir mounted on the lower breadboard. A pump (Amazon, DIPump-KK300-US) transferred media back to the top reservoir to maintain circulation. Each reservoir had a Millex-GS filter for oxygen exchange, and 5% CO_2_ was supplied to the bottom reservoir during imaging.

### Data correction files

Before imaging, field correction files were generated using the Cairn image Splitter calibration, a stack of 10 images of the NanoGrid acquired with transmitted light illumination. After imaging, flat-field corrections were acquired using two stacks of 10 images: one of fluorescein (ACROS ORGANICS, 2321-07-5), excited by the 488 nm laser, and one of DiD (Invitrogen, D7757), excited by the 647 nm laser. Stock fluorescein was prepared in 1 M NaOH at 1 mg/mL and diluted to 5 μL/mL in NaOH. DiD was diluted at 5 μL/mL in EtOH. The fluorophore solutions were added to Attofluor Cell Chambers (Invitrogen, A7816) for imaging. TIRF images of both coverslips were acquired with simultaneous excitation at 488 and 647 nm to mimic live-cell imaging conditions.

### STAR image processing

STAR Data was processed using DrSTAR in MATLAB 2025b. DrSTAR splits raw images into two channels: TIRF-EGFP and TIRF-iRFP713. For each channel, DrSTAR performs background subtraction, flat-field correction, bleach correction, interpolation correction, image registration, and pixel-by-pixel division of TIRF-EGFP and TIRF-iRFP images to generate Δz images, with Δz calculated relative to the local membrane.

### Tracking and analysis of clathrin dynamics

cmeAnalysis running in MATLAB 2025b was used to track CCSs output from DrSTAR. First, cmeAnalysis identified and tracked single diffraction-limited clathrin-coated structures (CCSs) based on the EGFP channel. The CCSs were divided into two groups based on the Δz signal, clathrin-coated vesicles (CCVs) and flat clathrin lattices (FCLs). To be classified as a CCV, the Δz signal rose above the baseline threshold for five consecutive frames, indicating the clathrin structure moved toward the cytosol. To be classified as a FCL, the Δz signal did not exceed the baseline threshold for five consecutive frames throughout the puncta lifetime, indicating this CCS did not change axial position. The total number of CCVs and FCLs identified in each cell was divided by the cell area and the imaging time (5 min), resulting in frequency per μm^2^ per min.

The vesicle formation model was determined for each individual CCV. The time clathrin was recruited was determined from the EGFP channel and the time the plasma membrane started curving was determined from the Δz channel. Both were identified as the first timepoint when the signal was above the baseline threshold for the next five contiguous frames (1.5 s).

### Statistics and reproducibility

All live-cell experiments were performed using three independent sets of transfections, unless stated otherwise. Statistical analyses were conducted in GraphPad Prism (Version 10.0.2).

## Funding

This work was supported by American Heart Association PRE906086 to T.N., American Heart Association PRE1191647 to J.Y., and NIH/NIGMS GM131099 to A.L.M.,

## Competing interests

The authors have no competing interests to declare.

## Author contributions

Conceptualization: JY, TJN, LDS, YTBT, and ALM; Investigation: JY; Methodology: JY, TJN, LDS, and YTBT; Writing – original draft: JY; Writing – review and editing: TJN, LDS, YTBT, and ALM; Funding acquisition: JY, TJN, and ALM.

---

**Supplemental Figure 1.**
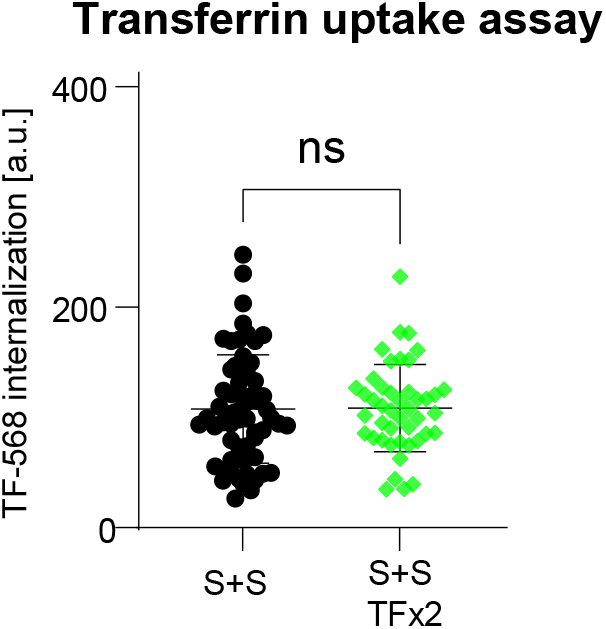
Tf uptake experiments were conducted at saturating concentration. HUVECs were cultured in static conditions and treated with 25 μg/mL (S+S) or 50 μg/mL of transferrin-AF568 (S+S TFx2) for 20 min. Plotted is the average transferrin fluorescence per cell for each group. S+S, 61 cells; S+S TFx2, 44 cells; from one independent experiment. Data analyzed by Welch’s test with mean ± SD presented.

## Notes

### Competing Interest Statement

The authors have declared no competing interest.

